# Molecular dynamics suggests antiviral compounds active against Dengue Virus show similar binding patterns to Zika Virus proteins

**DOI:** 10.1101/309351

**Authors:** Daniel Ferreira de Lima Neto, Anderson Pereira Soares, Shahab Zaki Pour, Ayda Susana Ortiz Baez, Patrick de Castro Neuhaus, Caio Cesar de Melo Freire, Carlos Francisco Sampaio Bonafé

## Abstract

The Zika virus (ZIKV) arrival in Brazilian territory brought to light the need for preparedness regarding arboviruses in Brazil. Compound screening is a cumbersome process dependent upon in vitro testing and validation. Recently, virtual screening methods have improved precision and reliability providing a framework for in silico testing of lead compound candidates. Here we have applied these methods on compounds that were previously shown to be active against Dengue virus in vitro, taking the structural information of such compounds and applying docking methods to identify putative binding sites. A molecular dynamics approach was also used to refine the docking results. The computational experiments ran here suggests that compounds such as Epigallocatechin Gallate, Ergotamine and Avermectin-B1a bind to active sites on the viral enzymes NS5 and NS3, as well as on its Envelope protein. Refinement shows that such bindings were not lost during the production run and key regions on both enzymes were structurally displaced on average over the simulation time. Interestingly there is no documented drug interactions among these candidates, raising the possibility of drug combinations during treatments. Moreover, the candidate compounds have been extensively studied, thus providing important information regarding intracellular interactions caused by them, which are also associated with pathways exploited by the virus, suggesting possible side interactions hindering the replication process.

## Introduction

Considering the main consequences of the ZIKV epidemics caused and the political and economical context the country, it became mandatory to search for antiviral compounds, a process which is often hindered by the overwhelming amount of experimentation necessary to test each candidate against the different settings needed to elevate the compound to a potential drug candidate status (Ghemtio et al., 2012; Veljkovic et al., 2007). The whole process is demanding in terms of the infrastructure required and the trial and error steps needed to produce lead compounds. The core facilities designed to iteratively screen FDA approved compound libraries against selected targets can be time-consuming in the event of a crisis such as the ZIKV epidemic (Saiz and Martín-Acebes, 2017).

The use of structure-activity relationships (SAR) has emerged as a suitable alternative for experimental compounds to be tested and has also been shown to reduce research time and also the financial cost invo0lved in drug discovery. A more accurate approach, termed Quantitative Structure-Activity Relationships (QSAR), consists in combining known functions of molecular compound features. In such methodology, activities of a given compound can be predicted quantitatively. Several models were made available over the years and are in use by major drug companies and research laboratories around the world (Putz et al., 2016; Richter et al., 2004; Verma et al., 2010).

Parallel to the developments of *in silico* drug testing, the computational structural biology groups also developed their methods to generate protein models based on its secondary and tertiary structures for downstream analysis and hypothesis testing. Through the combined information in public databases, ranging from sequencing to crystallographic experimentation, it has become commonplace to generate homology models for a designated target, provided that there is sufficient information in the databases to cover for secondary and tertiary structures coordinates (Agnihotri et al., 2012; Fernando and Fernando, 2017; Shiryaev et al., 2017). In addition, it is of the utmost importance that the genetic information about the viral strains to properly feed the model building process is also present.

## Methods

### Sequence analyses

Based on recent data from ZIKV virus sequences obtained from GenBank, complete genomes were selected to evaluate clusterization. Sequences chosen for this work were selected based on phylogenetic reconstructions in order to identify their distances from other documented sequences and to support their selection for homology modeling steps. Sequences from the strain circulating in Brazil were aligned with Muscle 3.8.31 (Edgar, 2004). Likewise, duplicates were removed using USEARCH v9.0.2132 (Edgar, 2010). Phylogenetic relationships were reconstructed by means of PhyML v3.0 software with 1000 bootstrap replicates (Guindon et al., 2005). The best-fit model of nucleotide substitution was selected in jModelTest v2.1.7 (Posada, 2008). The maximum likelihood tree was visualized and edited with FigTree v1.4.3.

### ADME/Tox predictions

Twenty-nine compounds from published articles with inferred mechanisms of action especially against DENV and the other flaviviruses were chosen. The structural data of these molecules were obtained from PubChem (http://pubchem.ncbi.nlm.nih.gov) and checked for errors and violations with the java based chemical drawing tool MarvinSketch (http://www.chemaxon.com). ADMET predictions were carried out with two different prediction tools, VEGA (Virtual evaluation of chemical properties and toxicity - https://www.vegahub.eu/) from QSAR models and Toxtree (Toxic Hazard Estimation by decision tree approach) (Patlewicz et al., 2008). Results were crosschecked with the SwissADME tool (Daina et al., 2017). Visualization of the chemical space used and principal component analysis (PCA) were run using DataWarrior (Sander et al., 2015).

### Homology modeling

Based on the phylogenetic clustering obtained for the chosen ZIKV sequences, corresponding crystallographic structures were downloaded from the Protein Data Bank as part of the homology modeling protocol implemented in the I-TASSER standalone software (Roy et al., 2010) to recreate models for the Brazilian sequences available at this point. Each complete genome was then broken into the amino acid sequences to be modeled. Conditions for homology modeling were as follows: 10 PSI-Blast iterations (E-Value of 0.1), twenty templates with five sequence alignments per template were used to build the hybrid model. Modeling was set to low speed with ten terminal extensions, sampling fifty terminal loops. Outputs were structurally aligned and compared for RMSD deviations with recent crystallized structures available for ZIKV proteins. Coordinate files obtained for the structural proteins (capsid, C; envelope, E and matrix, M) and for the non-structural proteins as well (NS1, NS2a, NS2b, NS3, NS4a, NS4b and NS5) were checked under PROCHECK (https://www.ebi.ac.uk/thornton-srv/software/PROCHECK/) for φ and Ψ violations. PDB coordinate files were then used in downstream analysis. Energy minimization of the 3D structures was performed using YASARA Structure, which runs molecular dynamics simulations of models in explicit solvent, using a new partly knowledge-based all atom force field derived from the force field Amber99SB (Land and Humble, 2018).

### Dockings

Ligand preparation was made with the package AnteChamber from the software AmberTools 18 (D.A. Case and P.A. Kollman, 2018), briefly each compound was submitted to charge correction inside the selected forcefield (AMBER99LB), missing parameters and corrections were saved for each compound and using the LeaP package inside Amber18, the corresponding topology files were corrected and converted to GROMACS (Abraham et al., 2015) topologies using the ACPYPE python script (Sousa da Silva and Vranken, 2012). For each ligand topology created this way a docking screening was made against each modeled protein using the VINA binary module, to compute the binding energy and the dissociation constants of the docked ensembles. Coordinate files of both ligand and receptors were submitted in PDBQT format with the generated coordinate and charge parameters after both structures had been corrected for missing atoms and energy minimized in 0.9% saline simulation box. Parameterization was kept as default without restricting docking to previously identified active sites. Twenty-five runs were briefly made for each pair ligand-receptor for all ZIKV modeled proteins, saving the lowest energy complex per cluster, which is based on the RMSD distance between each, set to 5.0 Å.

### Molecular Dynamics

The highest binders were analyzed as docked ensembles (ligand and receptors) and submitted to molecular dynamics (MD) production. Briefly, a dodecahedron simulation box was created around all atoms of the model. The simulations box size varied accordingly based on the proteins used. Missing atoms and parameters were corrected in a previous step. Water molecules were used to fill the box complemented with 0.9% of Na+ and Cl-ions to achieve electrostatic neutrality and the pH was set to physiological (7.4). Topology files previously created for the ligands were then inserted in the topology file of the protein of interest and proper modifications were made to run a ligand-receptor docking refinement using GROMACS. The AMBER99SB force field was used in periodic boundary conditions, temperature and pressure were kept at 300 K and 1 atm using long-range coulomb forces (Particle-Mesh Ewald). Production runs were allowed to run for 10 ns for the ligand-receptor complexes and also to each ZIKV modeled protein. Trajectory files were analyzed for structure RMSD, secondary structures, RMSD and RMSF calculations per residue. Average interactions were investigated using the software LigPlot and visualizations were prepared with the PyMol package. Non-docked proteins were used for comparison purposes and known antivirals against specific targets were used when applicable.

## Results

### ADME/Tox predictions

A literature review was made to select compounds with proven in vitro activity against Dengue Virus. Since DENV and ZIKV share a common ancestral, it was posited that the mechanisms of actions against the first could apply to the latter as well (Supplementary figure (SF1). These compounds characteristics are summarized in supplementary table 1, considering information on their putative mechanism of action, which experimental setting such claims were made on and the corresponding reference. Compounds that were not listed in PubChem were manually drawn using MarvingSketch tool, after which an energy minimization step was carried out and the structure was saved in compatible formats for downstream applications. Each compound is represented in its planar formula in figure 1 (FIGURE 1). The chemical space based on structure similarity is represented as a PCA graph (SF2) showing no overlapping structures selected for downstream analyses.

**Figure 1:**
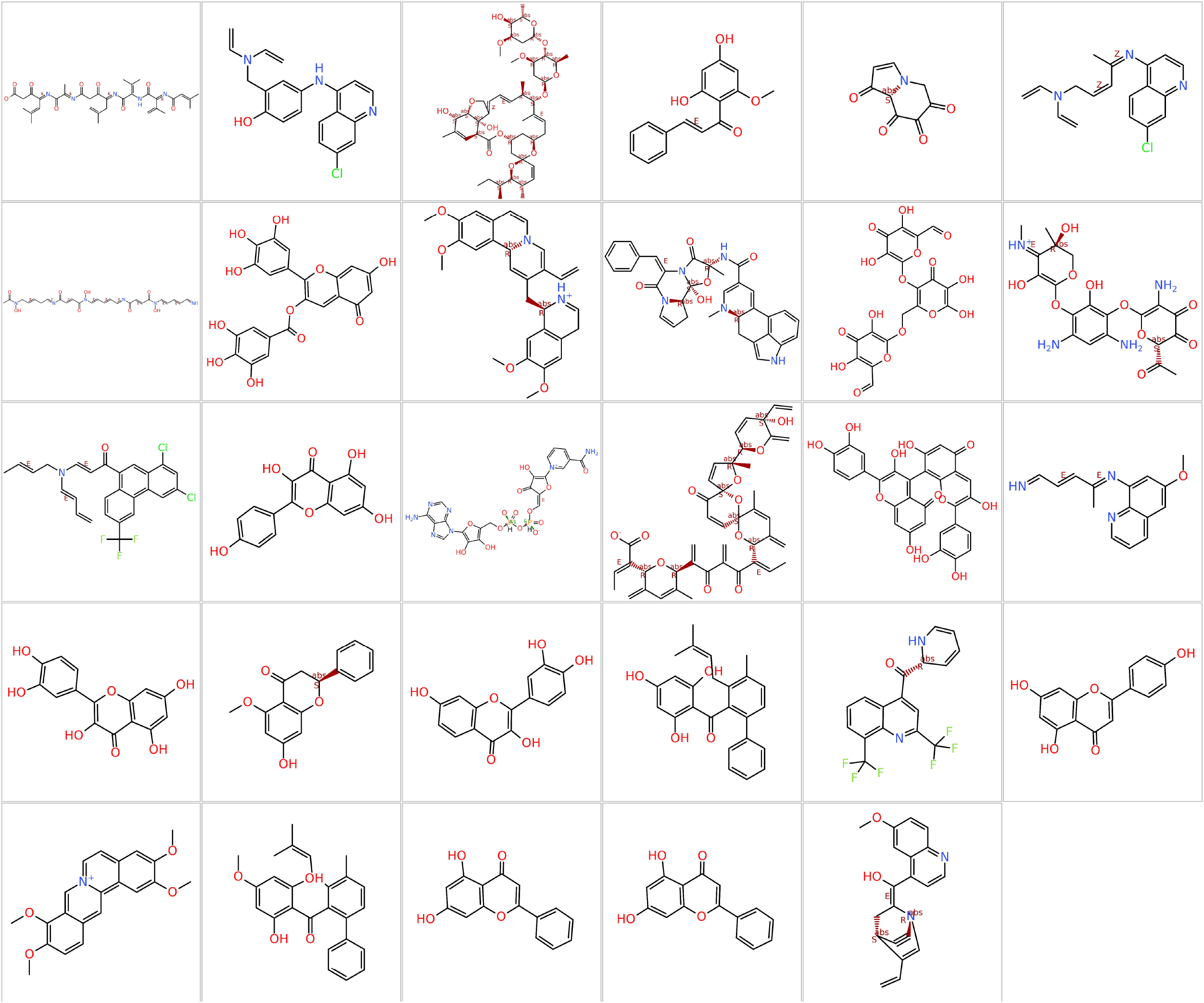
Compound dataset selected from the literature. A – Ahpatinin, B – Amodiaquine, C – Avermctin-B1a, D – Cardamonin, E – Castanospermin, F – Chloroquine, G – Deferoxamine, H – Epigallocatechin Gallate, I – Emetine, J – Ergotamine, K – Galactonnan, L – Geneticin, M – Halofantrine, N – Kaempferol, O – Nadide, P – Narasin, Q – Pepstatin, R–Primaquine, S – Quercitin, T – Quindine, U – Alpinetin, V – Fisetin, W – Hydroxypanduratin, X – Mefloquine, Y – Naringenin, Z – Palmatine, AA – Panduratin-A, AB – Pinocembrin, AC – Pinostrobin.

In view of the urgent need for alternatives to treat ZIKV infections, a compound screening must also evaluate the pharmacokinetic and pharmacodynamic properties of putative drugs. In this study, the selected compounds were evaluated for their absorption, distribution, metabolism, excretion and toxicological properties before the docking procedures. These results are summarized in supplementary table 1 (ST1).

Based on the results for this small dataset, it was possible to restrict the simulations could be restricted to three compounds (Ergotamine (ERGO), Avermectin 1Ba (AVE-1) and Epigalocatechin gallate EGCG)), considering their ADME/Tox properties, commercial availability and possible interactions within the host. Low absorption of these compounds was predicted, as opposed to the others in the dataset, but they are neither irritant nor tumorigenic, nor do they have effects on the reproductive tract, traits not shared by the other compounds taken together. A complete list of ADME/Tox properties is presented in supplementary table 1 (ST1).

### Homology modeling of ZIKV proteins

Based on the molecular epidemiology made available in the databanks, a phylogeny was recreated to guide the decision as to which strain proteins were to be modeled (SF1). The representative strain chosen is associated with the first ZIKV-associated microcephaly case sequenced and a complete genome was made available. Therefore, the open reading frames for the structural and non-structural proteins were selected and submitted to modeling. The procedure took advantage of recent findings in crystallography data produced for ZIKV as part of the structure selection process, generating closely related models directly related to the availability of crystallized ZIKV proteins, even though most were produced against the MR766 African strain PROCHECK and soft2 analysis ranked the modeled structures as optimal, with few residues in disallowed positions. The structures and the quality control tests for each model are represented in supplementary figure 3A to I.

### Compound docking to ZIKV proteins and molecular dynamics

Apart from the ADME/tox results, a virtual screening of the complete dataset was carried out against all modeled ZIKV proteins, considering previous reports of their in vitro antiviral activity. Our goal was to assess binding affinities and dissociation constants in the context of a productive viral infection, e.g. expressing the whole protein set. This result is presented in Figure 2 as a heatmap based on the binding affinities obtained after 25 docking rounds. Compounds such as Nadide (a dinucleotide or adenine and nicotinamide that has coenzyme activity in redox reactions and plays a role as a donor of ADP-ribose moieties) and Narasin (an antibacterial agent) docked with high binding affinity to key ZIKV proteins such as NS3 and NS5, but behave poorly in ADME/tox screenings. Conversely, the compounds ERGO, AVE-1 and EGCG had similar binding affinities to the same targets and were less toxic according to the in silico ADMET predictions. The binding affinities, Kd and docked regions’ raw results per protein are summarized in supplementary table 2 (ST2).

**Figure 2:**
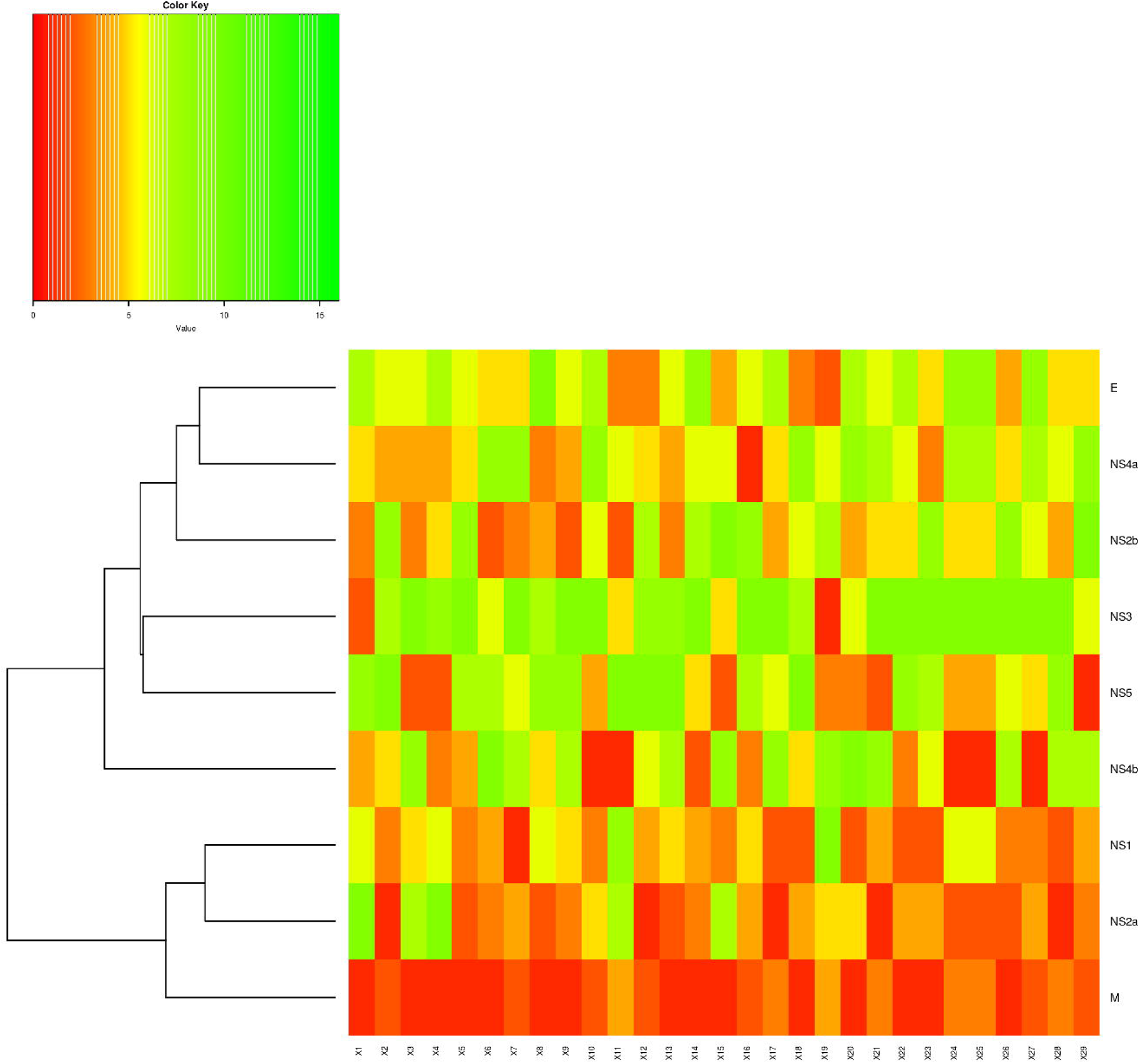
Heatmap of the virtual screening against all ZIKV modeled proteins. Higher binding affinities for each compound are represented as the transition from red to green. Columns represent each compound screened (X1 – X29) whilst lines represent each ZIKV protein.

The Envelope protein in Flavivirus plays key roles in the viral replication. It is subdivided into three domains, each responsible for a crucial step in the process. For this protein, the three selected compounds docked to approximately the same regions in domains I and III, varying in binding affinities and dissociation constants (ERGO > AVE-1 > EGCG - supplementary table 3 (ST3)). Even though no secondary structure rearrangements were detected over the 10 ns production run in comparison with the non-docked envelope (SF4a), significant RMSD and RMSF deviations were found in a per residue comparison. Whilst ERGO and AVE-1 dockings displaced residues in the vicinity of the fusion loop after the MD run (residues 80 – 120 (SF5a), fusion loop: residues 98 – 109), EGCG produced deviations in the 150 loop region of domain I, in both cases, the deviations were detected in the per residue RMSD and RMSF calculations. The best pose, average contacting residues and planar representations of the interactions are shown in supplementary figure 6 (SF6).

Another key player in the ZIKV replication events is the non-structural protein 3, NS3), comprising both the peptidase S7 function (residues 1 – 178), responsible for cleavage of the nascent polyprotein, the helicase ATP-binding domain (residues 181 – 337) and the helicase C-terminal domain (residues 332 – 511). The docking residues differed for each compound tested, as well as the binding affinities found (EGCG > ERGO > AVE-1 - supplementary table 3 (ST3)). The compounds interacted with residues from the ATP-binding domain of the helicase and only the ERGO docking suggests interactions with the peptidase region, associated wit the charge relay system for the serine protease activity. After the production runs for each condition, the overall behavior with regard to RMSF fluctuations of the ligand-receptor ensembles remained similar to the protein alone, except for the AVE-1:NS3 test, where RMSF per residues were markedly lower than the control, suggesting less mobility of this region in contrast with the other ensembles (SF7). Each pose and associated data (average contacting residues and planar representations) are shown in supplementary figure 7 (SF7). No significant secondary structure rearrangements were detected in this experiment either (SF4). On the whole, the data suggests that, if any inhibition in this system is to be experimentally found, the causes could be associated to the allosteric presence of the compound in functional sites, rather than to residue displacements causing loss of function.

Aside from its counterparts, the NS5 is the only target of a specific antiviral commercially available, thus providing a reasonable positive control to compare the selected compounds with. Without restricting the docking procedure to active sites, we were able to measure the highest binders and to compare these results against the documentation available for the drug Sofosbuvir. ERGO was the strongest binder, followed by EGCG and AVE-1, though different domains of the NS5 were targets for the compounds. EGCG and AVE-1 shared docking sites with the drug Sofosbuvir, while the compound ERGO docked against the initial sequence of the NS5, precisely to GTP-binding sites (ASN 17 and LYS 28). Other residues ERGO interacted with include the active sites of the methil transferase domain (residues 61, 146, 182 and 218). On the other hand, the residues interacting with EGCG and AVE-1 concentrated on the palm and thumb regions of the RNA dependent RNA polymerase domain of the NS5. The EGCG compound interacted with motifs A (motif: 532 to 543, docking: 535 to 539), E (motif: 709 to 715, docking: 712 to 713), and also with the priming loop (PL: 787 to 809, docking: 796 to 798) and the active site (AS: 664 to 666, docking: 665 to 666). AVE-1 interacted with the nuclear localization signal (NLS: 390), palm (479 to 708, docking: 495 to 525) and thumb (715 to 903, docking: 822 to 825), as shown in Figure 3. After the MD production, the perturbations caused by the ligands could be controlled, such as our positive control Sofosbuvir, these results are shown in supplementary figure 8 (SF8). The raw data for all simulations is available upon request. As expected, no significant secondary structure alterations were recorded along the simulation time (SF5).

**Figure 3:**
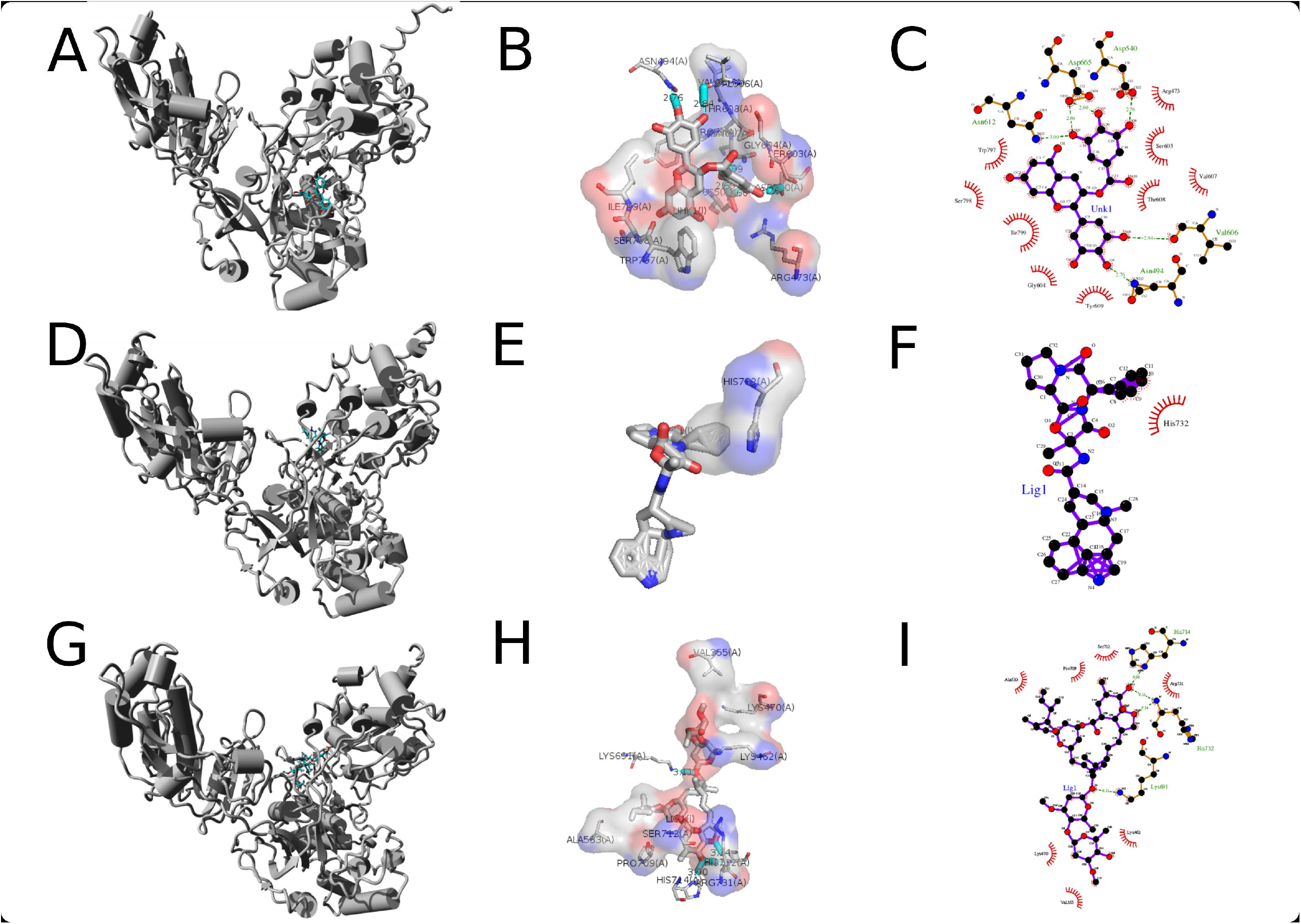
Docking refinement for the NS5 docked compounds-Poses and average residue interaction after a 10 ns molecular dynamics simulation for the top three screened compounds. A – NS5 – EGCG ensemble rendered in PyMol. B/C–EGCG interacting residues on average after MD (PyMol/LigPlot). D – NS5 – ERGO1 ensemble rendered in PyMol. E/F – ERGO1 interacting residues on average after MD (PyMol/LigPlot). G – NS5 – AVE1 ensemble rendered in PyMol. H/I – AVE1 interacting residues on average after MD (PyMol/LigPlot).

## Discussion

The use of in vitro settings to investigate and identify new compounds has made important advances over the past 100 years, particularly regarding the development of antivirals (Prusoff et al., 1989; Schinazi et al., 2009). More recently, new studies have relied on the ability to establish qualitative or semi-quantitative relations between molecular structures to test these hypotheses (Li, 2001). The computational (in silico) methods have been increasingly applied to virtual screenings, saving time and investments due to its ability to filter red flagged compounds (Dudek et al., 2006; Li, 2001 These in silico methods include databases, quantitative structure-activity relationships, pharmacophores, homology models and other molecular modeling approaches. Machine learning, data mining, network analysis tools and data analysis tools that use a computer are now seen as viable pathways for academic laboratories to compete with drug developers without major funding, saving time and resources and producing reliable results (Loregian and Palù, 2013). Screening for antivirals against ZIKV has been described as a race (Saiz and Martín-Acebes, 2017). Given the complications associated with the virus, the urgency is important and has produced several antiviral candidates at this point (Fernando and Fernando, 2017; Shiryaev et al., 2017), most of which are compounds repurposed to this end. Based on a literature search, we selected 29 compounds that were commercially available and had been previously shown to be active against Dengue (27) or Zika virus (2) and analyzed ADME/tox properties of this dataset to choose possible combinations to be used. We then asked, based on ADME/Tox predictions, which compounds could achieve key compartments without compromising important systems. We found that, in this small dataset, three compounds produced the best results, considering properties such as mutagenicity, tumorigenicity, druglikeliness, effects on the reproductive system and if it is an irritant, which produced a basis to question whether such compounds – previously shown active against Dengue in vitro – were capable of interfering with ZIKV as well. Considering that no structural alerts were detected and no protein nor DNA binding flags were raised, we asked if the putative antivirals had documented interactions with cellular proteins. Using the STITCH database (Szklarczyk et al., 2016) we found that ERGO1 maintains interactions with HTR1 family of genes. The protein encoded by this gene is a G-protein coupled receptor for serotonin (5-hydroxytryptamine) and it has been documented that ligand binding activates second messengers that inhibit the activity of adenylate cyclase and manage the release of serotonin, dopamine and acetylcholine in the brain. The compound is part of the family of alkaloids, being commonly used to treat migraines and severe headaches given its ability to cross the blood-brain barrier (Mulac et al., 2012). Depending on the administration route, effective levels of the drug could also be found in CSF of patients (Tfelt-Hansen et al., 2000). The drug currently is not recommended during pregnancy, possibly compromising its use by mothers during a productive ZIKV infection. Controlled studies have accessed effective dosage with minimal side effects in vivo (Silberstein and McCrory, 2003). The compound AVE1b and this macrolide family of anti-malaria drugs have been extensively tested against Flaviviruses and Alphaviruses (Mastrangelo et al., 2012) with impressive results regarding replication inhibition of these viruses. The study by Mastrangelo et al. analyzed the in silico and in vitro interactions the compound makes with its viral targets thoroughly and found that the putative mechanism of action is related to binding and interfering with the NS3 helicase activity. The ADME/Tox properties of the AVE1 compound have been extensively studied (Yang, 2012), making it possible to compare our in silico results with previously published studies of this compound. Taken together both in silico results and documented literature, these compounds present low toxicity and good levels of distribution, reaching effective concentrations against their original targets at the sites needed, e.g. ERGO1 interacts with BCRP/ABCG2, an ATP-binding cassette (ABC) efflux transporter involved in drug transport, which is highly expressed on the apical membranes of the placental syncytiotrophoblasts, the intestinal epithelium, the liver hepatocytes, the endothelial cells of brain microvessels, and the renal proximal tubular cells (Mao and Unadkat, 2015). Conversely, AVE1 has been found to induce autophagy in brain tissues on experimental exposure in pigeons (Wang et al., 2017) and Dou et al. have shown that the same class of antiparasitic drugs is associated with decreased P21-activated kinase 1 (PAK1) expression by modulating the ubiquitination-mediated degradation pathway (Dou et al., 2016, p. 1). The subsequent cascade lowers the phosphorylation levels of Akt, which finally results in the blockade of the Akt/mTOR signaling pathway, a pathway also altered in ZIKV infections (Chiramel and Best, 2017) EGCG on the other hand reaches high concentrations in plasma soon after administration (1.3 to 1.6h), produces harmless conjugates when excreted. Even though high levels interindividual variations in its pharmacokinetics profile were reported, the compound was previously linked to neurogenesis by several authors (Dube et al., 2011; Lee et al., 2002), and shown to be effective against ZIKV in vitro (Carneiro et al., 2016).

Considering the ADME/Tox characteristics of this small dataset, we then selected these compounds for docking screenings against the homology models of the ZIKV proteins, based on the genetic similarities both Dengue viruses and ZIKV share. We aimed to investigate previously reported antivirals against Dengue from an in silico structural perspective. At this point, a small number of high resolution crystallized structures of this virus are available, namely the NS2-NS3 complex and the polymerase NS5, and both crystals were based on the MR766 strain of the virus, which drove our efforts of homology modeling into using the Asian strain as a starting point. As a great number of crystallized structures are available for related Flavivirus, our approach was successful for most proteins, except for the NS4a and NS4b models, which ranked poorly in model evaluation protocols (SF3). These structures have few residues in disallowed regions, and were over 90% above the threshold for quality control, so that they could not be considered for downstream evaluations. Homology modeling has become an important tool in virtual screening reports (Agnihotri et al., 2012; Fernando and Fernando, 2017), and particularly useful in finding putative antivirals by molecular docking to homology models (Fernando and Fernando, 2017; Putz et al., 2016). Based on our models, we then sought to investigate possible interactions between the selected compounds and the viral proteins using an all versus all strategy and recording the docking positions as well as their dissociation constants. We found that binding affinities varied according to which protein a given compound was tested against, but our dataset – selected on the basis of previous reported activity against Dengue viruses of each compound – was therefore purposely biased against Flaviviruses. Comparing all modeled proteins with each docking result revealed that the selected compounds were bound to proteins that are key to viral replication, such as the envelope, the protease NS3 and the RNA polymerase RNA dependent NS5, giving support to our premise of shared activity against ZIKV and producing putative insights with regard to functional sites of the proteins the compounds were bound to. Furthermore, it also suggests possible interference mechanisms by which the virus replication process may be hindered, e.g. binding of the compounds to the priming loop of the NS5 RpRd or a putative blocking of the helicase activity of the NS3 protein, thus providing further insights into the action mechanisms of each compound against Dengue virus, as originally suggested by the respective authors of each original study used to populate our compound dataset, whilst introducing in silico data of their putative action against ZIKV as well.

As part of the screening procedures, we included a molecular dynamics step for both ligand-protein complexes and the protein alone for comparison purposes. This strategy has been applied before for membrane bound receptors as well as for viral proteins (Elfiky and Elshemey, 2018; Hornak et al., 2006). These steps were crucial to visualize perturbations across the protein structure as measured by RMSD/RMSF fluctuations over the simulations, in contrast with the ligand-free control for each case (SF6. SF7, SF8). Additionally, the use of a known antiviral drug as positive control for the NS5 MD experiments provided further insights into the similarities the selected compounds could share with it, as shown in Figure 3. Previous studies have suggested that the activity of enzymes is directly correlated with the flexibility of their active site, connecting rigidity with loss of function in most cases (Khan et al., 2016; Rashin et al., 2010). Ligand interactions with its target site increases side-chain rearrangements and may also contribute to conformational changes otherwise bound to enzymatic processes. The time scale associated with domain motions is observed in larger simulations, up to milliseconds or more. The association between structure fluctuations and enzymatic reactions can be investigated using MD simulations. Such experiments made on substrate-free and bound cyclophilin A (CypA) by McGowan provided important evidence as to the motions of active site residues in the complex, suggesting that the stabilization of the loop region is key to enzyme-substrate complex formation (McGowan and Hamelberg, 2013). These protein population shifts, due to conformational fluctuations derived from ligand binding, strengthens the intrinsic characteristics of the protein dynamics subject to conformational transitional states that can be stabilized by ligand binding (Vogt et al., 2014; Weikl and Paul, 2014), as proposed by our work on the interactions of ERGO1, AVE1 and EGCG with ZIKV key enzymes, as well as with the envelope protein (SF9, SF10 and F3).

Taken together, these simulations suggest plausible alternatives to address the ZIKV virus infections. Such claims need to be verified *in vitro*, a major limitation of our study. Even though precautions were taken to choose from compounds that were previously verified to be active against the Dengue viruses and given the close proximity DENV and ZIKV share phylogenetically, the gold standard still remains classical in vitro testing. The models, dockings and simulations herein are to be interpreted as a useful guide in further testing, hopefully narrowing down the possibilities to be tested. Conversely, there is considerable evidence to support these methods as fundamental tools to understand antiviral interactions with its putative targets. One can use the simulations results to build a cost-effective experimental framework, incorporating key aspects such as modeling of regions under selective pressure and the associated conformational changes to it. This creates a pipeline for drug screening that can be adapted to sequencing data and, therefore, account for important mutations RNA viruses undergo which can confer adaptive fitness to the virus. Integrating the abundance of sequencing data available with molecular epidemiology information on the viral targets under this framework is paramount to evaluate lead compounds.

---

**Supplementary figure 1**: Phylogenetic clustering of the reference ZIKV sequence (blue) used for homoology modeling.

**Supplementary figure 2**: Principal component analysis of the chemical space of the compounds selected for virtual screening considering druglikeliness parameters. Structure similarity is shown from red to blue.

**Supplementary figure 3**: Ramachandran plots of the modeled structures. A – Envelope, B – Capsid, C – NS1, D – NS2a, E – NS2b. F – NS3, G – NS4a, H – NS4b, I – NS5.

**Supplementary figure 4**: Secondary structure fluctuations during the molecular dynamics production run. Left – A – Envelope-EGCG ensemble, B - Envelope-ERGO1 ensemble, C - Envelope-AVE1-ensemble. Right – D – NS3 (ligand absent), E – NS3-EGCG ensemble, F - NS3-ERGO1 ensemble, G - NS3-AVE1 ensemble. Alpha helices are colored magenta, beta sheets are colored yellow, turns are colored pale blue, and all other residues are colored white

**Supplementary figure 5**: A – NS5 (ligand absent), B – NS5-EGCG ensemble, C – NS5-ERGO1 ensemble, D – NS5-AVE1 ensemble and E – NS5-Sofosbuvir ensemble. Alpha helices are colored magenta, beta sheets are colored yellow, turns are colored pale blue, and all other residues are colored white

**Supplementary figure 6**: RMSD and RMSF comparisons – Envelope trimer MD. Each docking refinement was made in parallel with the target protein without the ligand for control purposes. A – Per residue RMSD/RMSF plots of the subunit C of the trimeric envelope ensemble after the production run compared with the production run with the ligand AVE1b. B – Per residue RMSD/RMSF plots of the subunit C of the trimeric envelope ensemble after the production run, compared with the production run with the ligand EGCG, C – Per residue RMSD/RMSF plots of the subunit C of the trimeric envelope ensemble after the production run compared with the production run with the ligand ERGO1.

**Supplementary figure 7**: RMSD and RMSF comparisons – NS3 MD. Each docking refinement was made in parallel with the target protein without the ligand for control purposes. A – Per residue RMSD/RMSF plots of the NS3 ensemble after the production run, compared with the production run with the ligand AVE1b. B – Per residue RMSD/RMSF plots of the NS3 ensemble after the production run, compared with the production run with the ligand EGCG. C – Per residue RMSD/RMSF plots of the NS3 ensemble after the production run, compared with the production run with the ligand ERGO1.

**Supplementary figure 8**: RMSD and RMSF comparisons. A – Per residue RMSD/RMSF plots of the NS5 ensemble after the production run, compared with the production run with the ligand AVE1. B – Per residue RMSD/RMSF plots of the NS5 ensemble after the production run, compared with the production run with the ligand EGCG. C – Per residue RMSD/RMSF plots of the NS5 ensemble after the production run, compared with the production run with the ligand ERGO1. D – Per residue RMSD/RMSF plots of the NS5 ensemble after the production run, compared with the production run with the ligand Sofosbuvir.

**Supplementary figure 9**: Docking refinement for the Envelope docked compounds-Poses and average residue interaction after a 10 ns molecular dynamics simulation for the top three screened compounds. A – Subunit C (ENV)-EGCG ensemble rendered in PyMol. B/C–EGCG interacting residues on average after MD (PyMol/LigPlot). D – Subunit C (ENV)–ERGO1 ensemble rendered in PyMol. E/F – ERGO1 interacting residues on average after MD (PyMol/LigPlot). G – Subunit C (ENV)–AVE1 ensemble rendered in PyMol. H/I – AVE1 interacting residues on average after MD (PyMol/LigPlot).

**Supplementary figure 10**: Docking refinement for the NS3 docked compounds-Poses and average residue interaction after a 10 ns molecular dynamics simulation for the top three screened compounds. A – NS3–EGCG ensemble rendered in PyMol. B/C–EGCG interacting residues on average after MD (PyMol/LigPlot). D – NS3–ERGO1 ensemble rendered in PyMol. E/F – ERGO1 interacting residues on average after MD (PyMol/LigPlot). G – NS3–AVE1 ensemble rendered in PyMol. H/I – AVE1 interacting residues on average after MD (PyMol/LigPlot).

**Supplementary table 1**: Absortion, distribution, metabolism, excretion and toxicity properties of the compound dataset in use here. Lines correspond to the compound order presented in figure 1.

**Supplementary table 2**: Raw virtual screening results. Binding affinities and dissociation constants after 25 runs with VINA are presented with its corresponding docking receptor in columns.

